# Unveiling Multi-level and Multi-modal Semantic Representations in the Human Brain using Large Language Models

**DOI:** 10.1101/2024.02.06.579077

**Authors:** Yuko Nakagi, Takuya Matsuyama, Naoko Koide-Majima, Hiroto Q. Yamaguchi, Rieko Kubo, Shinji Nishimoto, Yu Takagi

## Abstract

In recent studies, researchers have used large language models (LLMs) to explore semantic representations in the brain; however, they have typically assessed different levels of semantic content, such as speech, objects, and stories, separately. In this study, we recorded brain activity using functional magnetic resonance imaging (fMRI) while participants viewed 8.3 hours of dramas and movies. We annotated these stimuli at multiple semantic levels, which enabled us to extract latent representations of LLMs for this content. Our findings demonstrate that LLMs predict human brain activity more accurately than traditional language models, particularly for complex background stories. Furthermore, we identify distinct brain regions associated with different semantic representations, including multi-modal vision-semantic representations, which highlights the importance of modeling multi-level and multi-modal semantic representations simultaneously. We will make our fMRI dataset publicly available to facilitate further research on aligning LLMs with human brain function. Please check out our webpage at https://sites.google.com/view/llm-and-brain/.

## 1 Introduction

Language models, which learn the statistical structure of languages from large corpora to enable machines to understand semantics, have made significant advances (Devlin et al., 2018; Radford et al., 2019; Zhang et al., 2022; Touvron et al., 2023). Because semantic comprehension is fundamental to human intelligence, the correspondence between human brain activity and the latent representations of language models has been an intriguing subject of research. To examine the correspondence between human and language models, in recent studies, researchers used **brain encoding models** (Naselaris et al., 2011; Nishimoto et al., 2011; Huth et al., 2012) to predict brain activity based on the high-dimensional latent representations of language models (Jain and Huth, 2018; Jat et al., 2019; Toneva and Wehbe, 2019; Caucheteux et al., 2021; Schrimpf et al., 2021; Goldstein et al., 2022; Caucheteux et al., 2022; Antonello et al., 2023). The quantitative assessment of the correspondence between the two can serve as a unique benchmark for modern large language models (LLMs) because it potentially provides a biological benchmark for the alignment between LLMs and humans.

In previous studies, researchers typically focused on a single aspect of semantic comprehension (e.g., speech content); however, realistic scenarios are inherently multifaceted. For instance, a scene in which two people are speaking can be depicted through multiple semantic levels in language: their speech content, their identities and the visual appearance of their outfits, the location and time of the scene, and the broader context of the conversation. In traditional studies, **researchers often address these levels of semantics separately**, which leads to a lack of clarity on how multiple levels of semantic content are attributed to brain activity and how the latent representations of language models might align with the human brain processing of multiple levels of content.

In this study, we address these issues by recording human brain activity using functional magnetic resonance imaging (fMRI) while participants view 8.3 hours of videos of dramas or movies. Importantly, we heavily annotate these videos across multiple levels of semantic content related to the drama, including speech dialogue, visual objects, and background story. We extract latent representations of these annotations from LLMs, then build encoding models that predict brain activity from these latent representations to quantitatively compare how each type of information is represented in different brain regions. Furthermore, **we quantitatively assess how each brain region uniquely captures different aspects of semantic content** using different types of latent representations derived from LLMs and multi-modal LLMs.

Our contributions are as follows:

1. Unlike previous researchers, who modeled various semantic modalities independently, we demonstrate that different semantic modalities uniquely account for brain activity in distinct brain regions.
2. We show that the superiority of LLMs is not uniform across different modalities: LLMs are particularly effective in modeling story-related information.
3. We show that latent representations of multi-modal vision-semantic LLMs predict brain activity and uniquely capture representations in the association cortex better than unimodal models combined.
4. We collect densely annotated fMRI datasets acquired while the participants watch 8.3 hours of videos as another benchmark for the biological metric of the alignment between LLMs and humans. We will publish these datasets.^1^

## 2 Related work

In numerous previous studies, researchers examined the relationship between the latent representations of language models and brain activity during speech comprehension. These researchers primarily focused on the correspondence between latent representations of language models obtained from speech transcriptions and brain activity (Huth et al., 2016; Jat et al., 2019; Toneva and Wehbe, 2019; Schmitt et al., 2021; Caucheteux et al., 2021, 2022). Recent findings have further demonstrated that LLMs provide a better explanation of brain activity than traditional language models (Schrimpf et al., 2021; Goldstein et al., 2022; Antonello et al., 2023; Tuckute et al., 2024).

Regarding the semantic content of the visual object in the scene, the correspondence between the latent representations of deep learning models related to the displayed objects and human brain activity have been studied extensively (Güçlü and van Gerven, 2015; Horikawa and Kamitani, 2017; Groen et al., 2018; Wen et al., 2018; Khosla et al., 2021; Allen et al., 2022; Chen et al., 2023; Wang et al., 2023; St-Yves et al., 2023; Takagi and Nishimoto, 2023; Luo et al., 2023). In most of these studies, researchers used neuroimaging data while participants watched static images or short video clips.

Although several neuroscience studies have been conducted in which researchers explored high-level story content in the brain using a naturalistic movie watching experiment (Hasson et al., 2008; Wehbe et al., 2014; Aw and Toneva, 2022; Chang et al., 2021; Nastase et al., 2021), these researchers typically did not explicitly model high-level story content using language models. Furthermore, they have not examined the unique explanatory power story-specific semantic content exerts on brain activity compared with other types of semantic content.

There is a growing consensus that LLMs mirror human brain activity more accurately than traditional language models during semantic comprehension; however, in the individual studies described above, the reseachers addressed discrete aspects of semantic comprehension independently. This is problematic because humans process different types of semantic content simultaneously in naturalistic scenarios, and such content may be represented differently in LLMs and the human brain. Here, we address these issues by evaluating how much each level of semantic content uniquely explains brain activity compared with other semantic content.

## 3 Methods

### 3.1 fMRI experiments

We collect brain activity data from six healthy participants with normal and corrected-normal vision (three females; age 22–40, mean = 28.7) while they freely watch 8.3 hours of videos of movies or drama series. All participants are right-handed, native Japanese speakers. They provided written informed consent for the study and the release of their data. The ethics and safety committees approved the experimental protocol.

MRI data are acquired using a 3T MAGNETOM Vida scanner (Siemens, Germany) with a standard Siemens 64-channel volume coil. Blood oxygenation level-dependent (BOLD) images are acquired using a multiband gradient echo-planar imaging sequence (Moeller et al., 2010) (TR = 1,000 ms, TE = 30 ms, flip angle = 60°, voxel size = 2×2×2 mm^3^, matrix size = 96×96, 72 slices with a thickness of 2 mm, slice gap 0 mm, FOV = 192×192 mm^2^, bandwidth 1736Hz/pixel, partial Fourier 6/8, multiband acceleration factor 6). Anatomical data were collected using the same 3T scanner using T1-weighted MPRAGE (TR = 2530 ms, TE = 3.26 ms, flip angle = 9°, voxel size = 1×1×1 mm^3^, FOV = 256×256 mm^2^). The preprocessing of the functional data includes motion correction, coregistration, and detrending. See Section A.1 for details of the acquisition and preprocessing procedures.

### 3.2 Stimuli

We use nine videos of movies or drama series as stimuli (10 episodes in total), with dense annotations related to those videos. The videos encompass various genres. Eight are international videos and one is a Japanese animation. The average play-back time of the 10 episodes is 49.98 minutes (ranging from 21 minutes to 125 minutes). We divide each episode into two to nine parts, each approximately 10 minutes long, for use as stimulus videos during the fMRI experiment. We play all the international videos in Japanese dubbed versions and the subjects understand them in Japanese. We will make our fMRI dataset publicly available for future research on acceptance. See Section A.1.1 for details of data collection.

The annotations include five levels of semantic content from the videos: transcriptions of spoken dialogue (*Speech*), objects in the scene (*Object*), background story of the scene (*Story*), summary of the story (*Summary*), and information about time and location (*TimePlace*) (Figure 1a). The semantic content consists of annotations that describe the stimulus videos in natural language. This content differs in the nature of the description and the timespan of the annotations. Specifically, *Speech* corresponds to the intervals of speaking, whereas *Object* is annotated every second, *Story* every 5 seconds, *Summary* approximately every 1-3 minutes, and *TimePlace* every time the screen changes. While there may still be some overlap among annotations, the motivation behind this annotation scheme is to capture different aspects of objective and narrative information. Multiple annotators independently label each level of semantic content, except for *Speech*: five for *Object*, three each for *Summary* and *TimePlace*, and two for *Story*. Before starting the work, annotators were explicitly informed that these descriptions are distinct from other annotations. For all five classes of semantic annotations, at least two researchers independently and regularly review the annotations, making corrections if any errors are found in the descriptions. While all annotations are originally described in Japanese, they are translated into English using DeepL and we used language models mainly trained on a English corpus.^2^ We also quantitatively confirm that the results are robustly reproduced when the annotators or data are divided. See Sections A.2 and A.3 for details of annotations and quality control.

**Figure 1:**
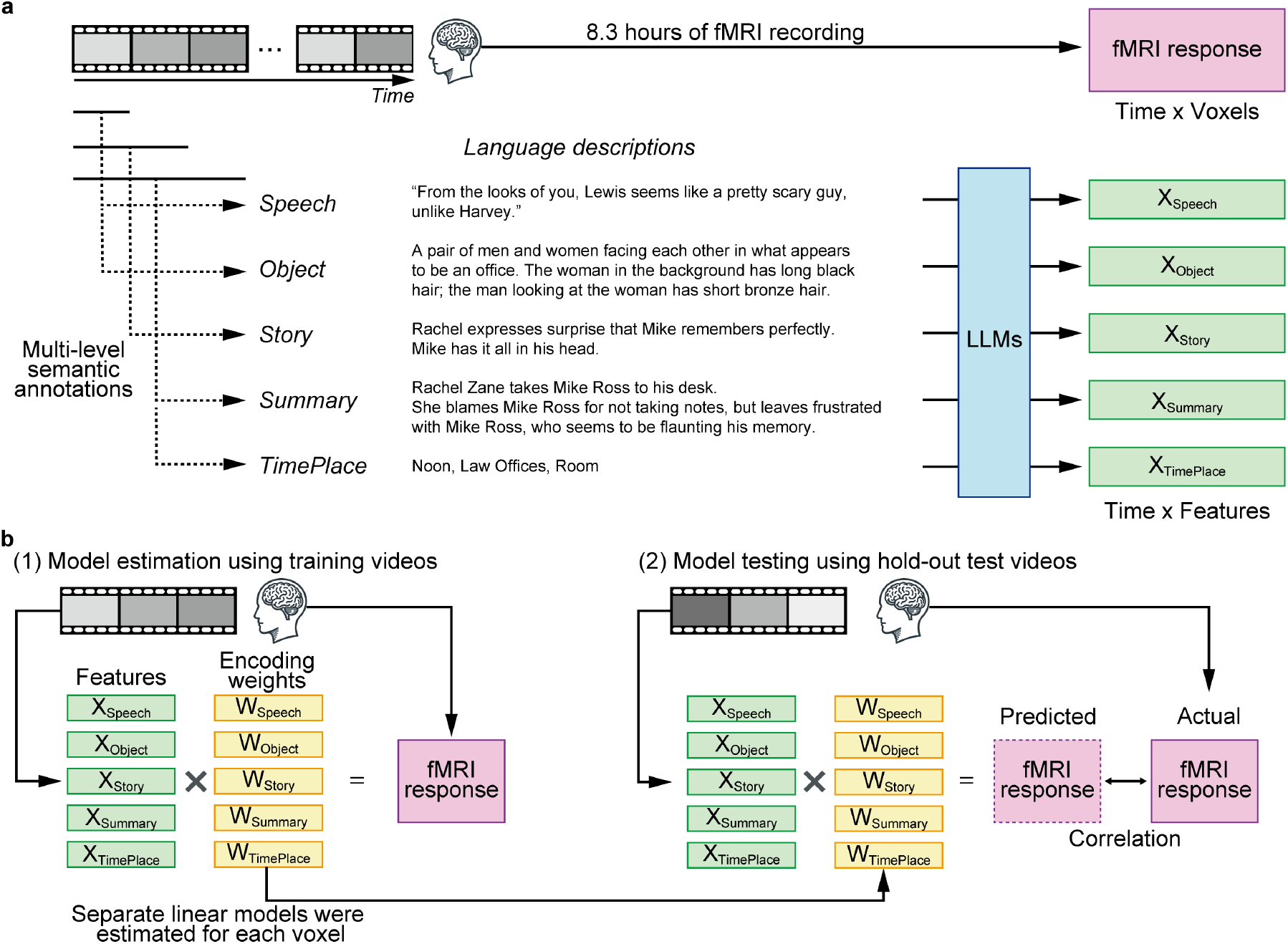
Overview of the experiment and brain encoding models. **a** In the experiment, participants watch 8.3 hours of videos of dramas or movies while we measure their brain activity inside an fMRI scanner. We densely annotate the videos with multiple levels of semantic content, which are used for extracting latent representations from multiple language models. The annotations provide examples from specific scenes in the series ‘*Suits*’. **b** For each semantic feature obtained by the language models, we estimate linear weights for predicting brain activity across the cerebral cortex from the feature using ridge regression. We subsequently apply the estimated weights to the features of the test data to estimate brain activity. We calculate prediction performance using Pearson’s correlation coefficient between the predicted and actual fMRI responses.

We divide the data into training and test datasets, and calculate all the prediction performance results presented in this paper using the test dataset. Specifically, we use the fMRI scanning sessions corresponding to the last split of each movie or drama series, 7,737 seconds in total, as test data. We use the remaining sessions, 22,262 seconds in total, as training data.

### 3.3 Feature extraction

We obtain latent representations from five language models (Figure 1a): Word2Vec (Mikolov et al., 2013), BERT (Devlin et al., 2018), GPT2 (Radford et al., 2019), OPT (Zhang et al., 2022), and Llama 2 (Touvron et al., 2023). We use Word2vec as a traditional language model. The summary of these language models is presented in Table 1. See Section A.1.5 for the information of the models we used.

**Table 1:** Summary of the five language models.

| Model | Layers | Width | Params. |
| --- | --- | --- | --- |
| Word2Vec<br>(Mikolov et al., 2013) | 1 | 300 | 900M<sup>3</sup> |
| BERT<br>(Devlin et al., 2018) | 12 | 768 | 110M |
| GPT2<br>(Radford et al., 2019) | 36 | 1280 | 812M |
| OPT<br>(Zhang et al., 2022) | 32 | 4096 | 6.7B |
| Llama 2<br>(Touvron et al., 2023) | 32 | 4096 | 6.74B |

We extract the latent representations of the language models for the annotations from each of the hidden layers, except for Word2Vec, from which we obtain word embeddings. For each latent representation from each hidden layer, we build brain encoding model (see Section 3.4). We first extract the latent representations of annotations, each of which consists of several tokens or words, for each time point. Then, we average the latent representations of the LLMs across tokens or average the word embeddings of Word2Vec across words. Because multiple annotations exist for each second, except for *Speech* annotations, we calculate the latent representations for each annotator for each language model and then average the latent representations across all annotators. For OPT and Llama 2, we reduce the dimensions of the flattened stimulus features using principal component analysis (PCA) and set the number of dimensions to 1280 (the same dimension as GPT2). We calculate the PCA loadings based on the training data and apply these loadings to the test data. In the main analysis we calculate latent representations using the annotation that correspond to the TR (1 second). We confirm that the results do not change significantly when we use longer context-lengths (see Figure A.8).

We also explore how uniquely different levels of semantic content explain brain activity compared with features from other modalities: vision, audio, and vision-semantic. For visual features, we extract latent representations from DeiT (Touvron et al., 2021) and ResNet (He et al., 2016). We input the first and middle frames of each second of the video into the model and extract the output from each hidden layer. For audio features, we extract the latent representations of audio data in the video using AST (Gong et al., 2021) and MMS (Pratap et al., 2023). We input audio data into the model in 1-second intervals. We extract the output from each hidden layer in response to the input. For both the vision and audio modalities, we reduce the dimensions of the flattened stimulus features using PCA and set the number of dimensions to 1280. We calculate the PCA loadings based on the training data and apply these loadings to the test data.

In addition to the unimodal feature, we extract vision-semantic multi-modal representations from GIT (Wang et al., 2022), BridgeTower (Xu et al., 2023), and LLaVA-v1.5 (Liu et al., 2023). To obtain vision-semantic representations, we fed paired visual and semantic data into the vision-semantic models, using the same input format as for the unimodal models. We use output from the text-decoder in GIT, the cross-modal encoder in BridgeTower, and the text-decoder in LLaVA-v1.5 as vision-semantic features. We average the latent representations of text and images across tokens, respectively. We finally reduce the dimensions of the flattened features using PCA and set the number of dimensions to 1280. In variance partitioning analysis used to investigate unique variance explained by multi-modal features compared with unimodal features (see Section 3.7), we use Vicuna-v1.5 (Zheng et al., 2024), which is the base model of the text decoder of LLaVA-v1.5, and CLIP (Radford et al., 2021), which is the image encoder of LLaVA-v1.5. For CLIP, we use the model in LLaVA-v1.5, and for Vicuna-v1.5, we use the model provided by lmsys.

### 3.4 Brain encoding models

To investigate how different levels of semantic content are represented differently in the human brain, we first build brain encoding models to predict brain activity from the latent representations of language models for each semantic content independently (see Figure 1b). We separately construct encoding models for each subject, feature, and layer (if applicable).

We model the mapping between the stimulus features and brain responses using a linear model *Y* = *XW*, where *Y* denotes the brain activity of voxels from fMRI data, *X* denotes the corresponding stimulus features, and *W* denotes the linear weights on the features for each voxel. We estimate the model weights from training data using L2-regularized linear regression, which we subsequently apply to test data. We explore regularization parameters during training for each voxel using cross-validation procedure. Because the dataset contains nine dramas and movies, to tune the regularization parameters during training, we select sessions from two to three dramas or movies as validation data and use the remaining videos as training data. We repeat this procedure across all dramas and movies. For the evaluation, we use Pearson’s correlation coefficients between the predicted and measured fMRI signals. We compute the statistical significance using blockwise permutation testing. Specifically, to generate a null distribution, we shuffle the voxel’s actual response time course before calculating Pearson’s correlation between the predicted response time course and the permuted response time course. During this process, we shuffle the actual response time course in blocks of 10TRs to preserve the temporal correlation between slices. We identify voxels that have scores significantly higher than those expected by chance in the null distribution. We set the threshold for statistical significance to *P <* 0.05 and correct for multiple comparisons using the FDR procedure. We conduct all encoding analyses using the *himalaya* library^4^(la Tour et al., 2022). We will make our code publicly available on acceptance. In the analysis for comparing different language models (Figures 2 and 3), we assume hemodynamic delays of 8-10 seconds from neural activity to the BOLD signal. We confirm that choice of delay time does not significantly affect on the results (See Figure A.7). In the analysis for comparing multi-modal features (Figure 5), we use the delay time of 6-8 seconds for the analysis of all features.

**Figure 2:**
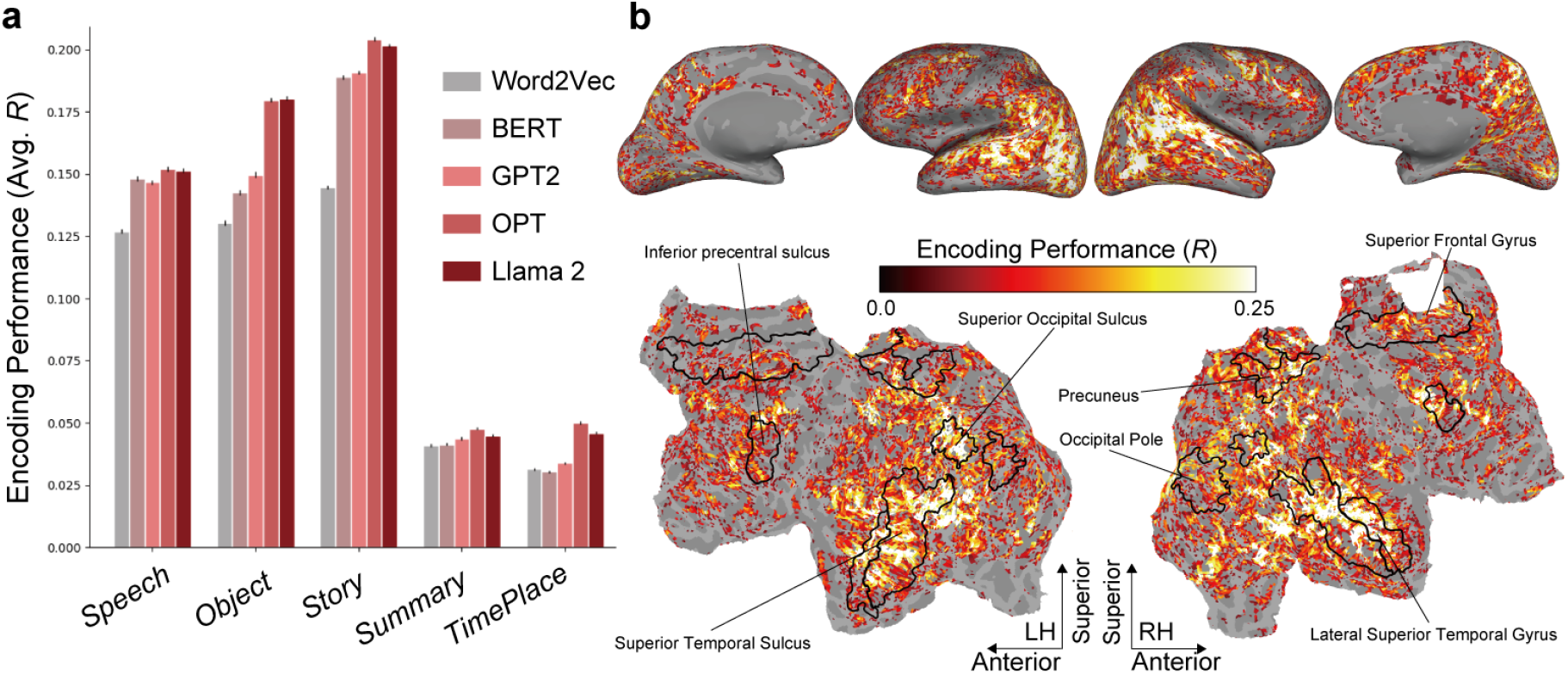
Encoding model results. **a** Prediction performance (measured using Pearson’s correlation coefficients) when predicting the brain activity of a single participant (DM09) from the latent representations of different language models for five distinct levels of semantic content. The figure presents the average prediction performance on the test dataset for the top 5,000 voxels, which we select within the training cross-validation folds in the layer that exhibits the highest prediction performance. We choose layers and voxels for each semantic content and each language model, respectively. The error bars indicate the standard error of the mean across voxels. **b** Prediction performance for a single subject (DM09) when all five levels of semantic content are used simultaneously using Llama 2, projected onto the inflated (top, lateral, and medial views) and flattened cortical surface (bottom, occipital areas are at the center), for both the left and right hemispheres. Brain regions with significant accuracy are colored (all colored voxels *P <* 0.05, FDR corrected). Black contours show several representative, anatomically defined regions of interest (ROIs)

**Figure 3:**
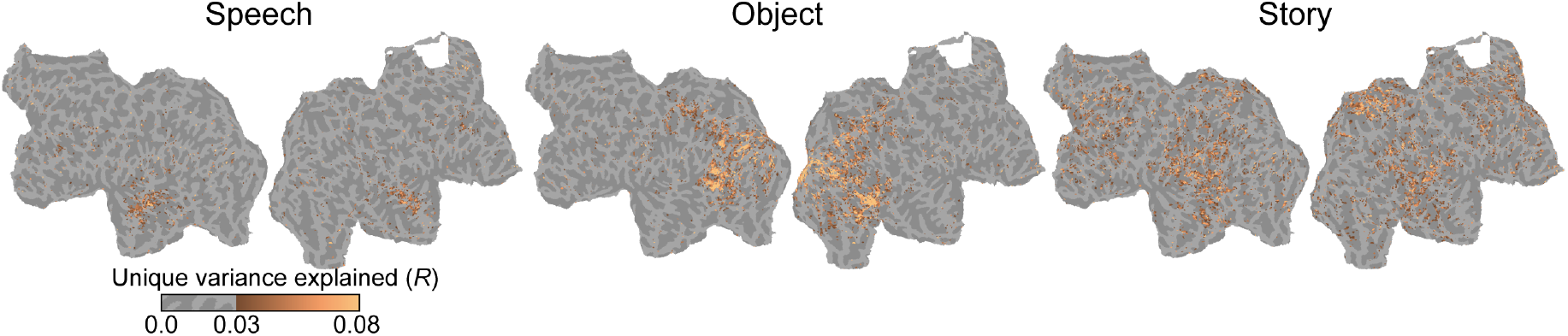
Variance partitioning analysis. Unique variance explained by the latent representations of the semantic content of (**a**) *Speech*, (**b**) *Object*, and **c** *Story* for a single subject (DM09). We use Llama 2 as a language model. For illustration purposes, we color only the voxels with a unique variance above 0.03.

### 3.5 Comparison of different levels of semantic content

To examine how uniquely each level of semantic content explains brain activity, we construct a brain encoding model that incorporates all the semantic features and evaluate the unique variance explained by each semantic feature using variance partitioning analysis (la Tour et al., 2022). In variance partitioning analysis, we determine the unique variance explained by subtracting the prediction performance of a model with a certain feature of interest removed from the prediction performance of the full model that includes all features. To estimate the model weights, we use banded ridge regression (la Tour et al., 2022), which can optimally estimate the regularization parameter for different feature spaces. For variance partitioning analysis, we use Llama 2 as LLMs, and use the latent representations from the layers that demonstrate the highest accuracy in cross-validation within the training data for each level of semantic content in the previous encoding model analysis.

### 3.6 Principal component analysis

For interpretation purposes, we apply PCA to the weight matrix of the encoding model (Huth et al., 2012). Here, we focus on the representation of the *Story* feature because the representation of such high-level semantic content in the brain has not been quantitatively evaluated in previous studies. We use only the voxels with top 5,000 prediction performance. To interpret the estimated PCs, we project randomly selected 1,640 *Story* annotations onto each PC, thus acquiring PC scores for the annotation. To further interpret the PC scores, we use GPT-4 (*gpt-4-1106-preview* in the OpenAI API) to classify these annotations into five semantic attributes that are commonly present throughout the annotations. Finally, we interpret PC1, PC2 and PC3 as axes that represent content related to the environment, interaction and cooperation in the drama respectively. See Section A.1.4 for details.

### 3.7 Comparison of different modalities

Taking advantage of the fact that our stimuli consist of visual, auditory, and semantic multi-modal elements, we compare the prediction performance of the visual, auditory, and semantic modalities of brain activity. Thus, we use not only unimodal features but also multi-modal features of the vision-semantic modality. We build the encoding model following the same procedure described previously and compare its whole-brain prediction performance.

Furthermore, we test whether the multi-modal features can predict brain activity components that unimodal features cannot. For this purpose, we use variance partitioning analysis as described earlier. The analysis procedure remains nearly identical to the procedure we use for the different levels of semantic content. We concentrate this analysis on LLaVA-v1.5, which performs particularly well among multi-modal models. As unimodal models, we use Vicuna-v1.5 for the semantic modality, CLIP for the vision modality, and AST for the audio modality. Here, we use *Speech* as semantic features.

## 4 Results

### 4.1 Comparison of the language models

We first evaluate how different levels of semantic content explain brain activity independently. Figure 2a shows that *Speech, Object*, and *Story* content predict brain activity with higher accuracy than *Summary* and *TimePlace* content. Notably, the larger the model, the better the prediction performance, and in particular, Llama 2 consistently achieves higher prediction performance than Word2Vec for all subjects for *Speech, Object*, and *Story* (*P <* 0.05, paired t-test). It also shows that large models achieve higher prediction performance for high-level background *Story* content. See Figure A.1 for the results for all subjects. Figure 2b shows the prediction performance of the whole-brain voxel-wise encoding model with all five levels of semantic content simultaneously with Llama 2. It demonstrates that we can predict brain activity across a wide range of brain regions involved in high-level cognition, in addition to sensory areas that include vision and audio.

### 4.2 Variance partitioning analysis

In the previous analysis, we showed that the LLMs’ latent representations of *Speech, Object*, and *Story* content predict brain activity well. However, the unique contribution of each level of semantic content to the explanation of brain activity remains unclear. Next, we use variance partitioning analysis (la Tour et al., 2022) to determine the extent to which the different types of semantic content uniquely account for brain activity.

Figure 3 shows that the latent representations of Llama 2 for the semantic content of *Speech, Object*, and *Story* correspond to spatially distinct brain regions. By contrast, *Summary* and *Time-Place* do not have unique variance (see Figure A.3). Specifically, *Speech* is associated with the auditory cortex, *Object* with the visual cortex, and *Story* with a broader brain region, including the higher visual cortex, precuneus, and frontal cortex. See Figure A.2 for the results at the level of ROIs. Furthermore, when we perform similar analysis using Word2Vec with *Story*, the unique variance is lower than that of Llama 2. This suggests that our encoding results obtained by LLMs reflect high-level semantic information representations in the brain (see Figure A.4).

These findings are consistent with those of previous researchers who focused on individual modali-ties (e.g. (Huth et al., 2016)). However, **a critical distinction from earlier work is that we focus on the unique explanatory power of individual modalities when compared with other modalities**; that is, building on insights from previous studies, we present the first comprehensive results that integrate various semantic content into a single study. Figure A.3 shows the results for all subjects for all semantic content.

### 4.3 Principal component analysis

Thus far, we have demonstrated that different levels of semantic content uniquely explain spatially distinct brain regions. Next, we analyze what specific information is captured by our encoding models by applying PCA to the weight of the encoding model for the latent representations of Llama 2 for *Story* content. The first three PCs explain the weight matrices of the top 5000 voxels, based on prediction performance, with explained variance ratios of 27.2 ± 4.5%, 13.7 ± 1.3%, and 7.6 ± 1.6% (mean ± s.t.d, N=6). Figure 4a presents the projection of PC scores computed for each voxel for two example participants (DM03 and DM09). While high-level semantic content within *Story* annotations is presumed to vary significantly between individuals, we observe certain trends across participants, particularly PC1 and PC2. See Figure A.5 for the results for all subjects.

**Figure 4:**
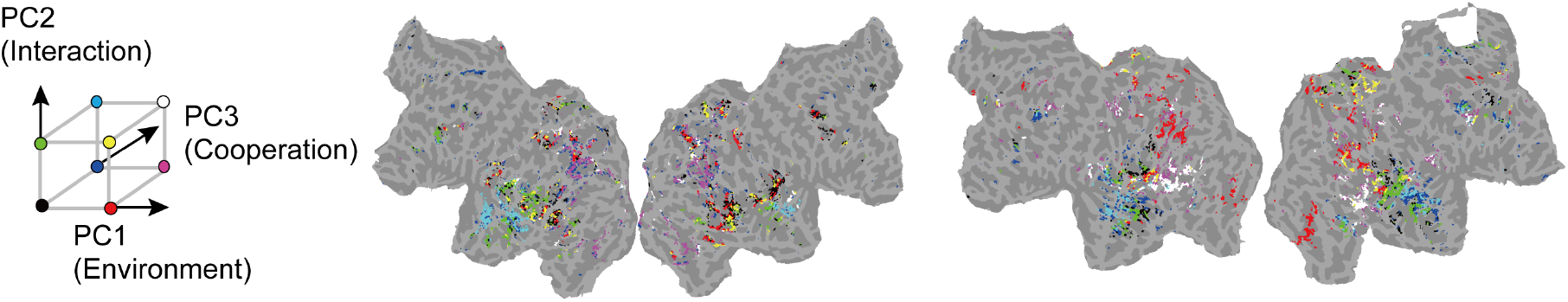
Principal component analysis. PCA on the encoding weight matrix of the latent representations of Llama 2 identifies the first three PCs for two subjects (Left, DM03; Right, DM09). Only the voxels with top 5,000 prediction performance are used for PCA.

### 4.4 Comparisons among the modalities

Given the multi-modal nature of the stimuli in our study, we can quantitatively compare the prediction performance of latent representations of semantics with that of other modalities, such as vision and audio.

Figure 5a shows the prediction performance of each modality’s latent representation. Similar to the semantic features, the visual and auditory features predict brain activity well. Interestingly, the multi-modal features (e.g. LLaVA-v1.5) predict brain activity better than all other features.

**Figure 5:**
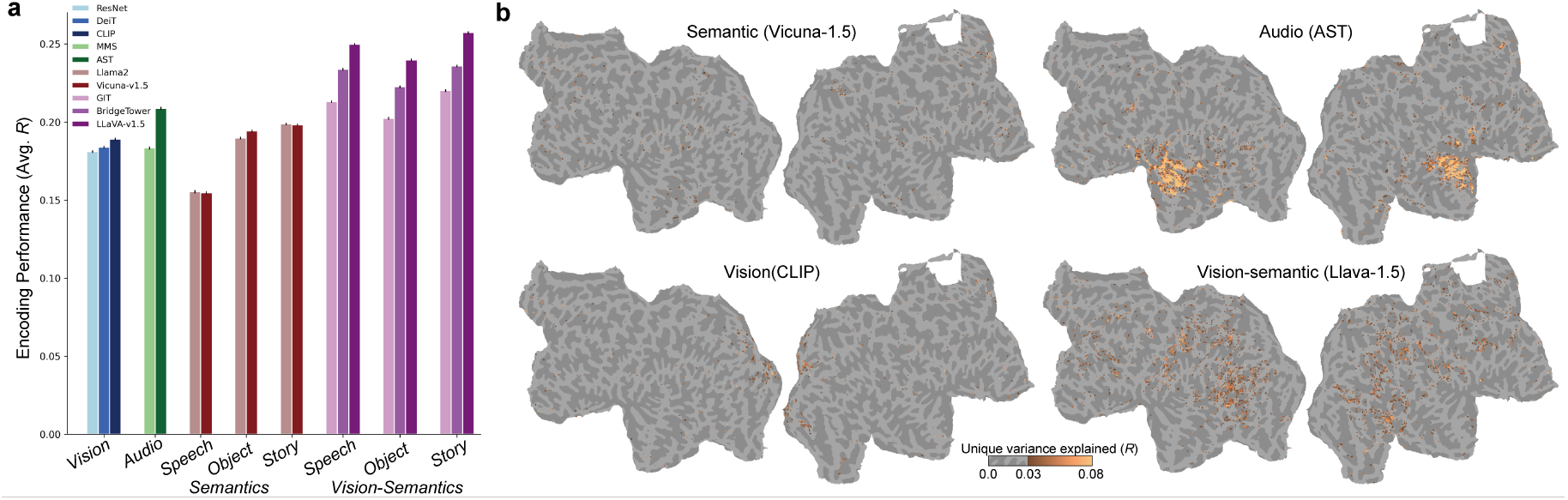
Comparisons among the modalities. **a** Prediction performance when predicting the brain activity of a single participant (DM09) from the latent representations of the feature modalities. The results for Llama 2 are the same as those in Figure 2b. **b** Unique variance explained by four modality features for a single subject (DM09). Here, we use *Speech* as semantic features for Vicuna-v1.5 and LLaVA-v1.5. For illustration purposes, we color only the voxels with unique variance above 0.03.

Figure 5b demonstrates that the multi-modal features (LLaVA-v1.5) uniquely predict brain activity in the association cortex, which cannot be explained by the unimodal features.

Together, these results suggest that the information processing style of state-of-the-art multi-modal deep learning models corresponds to human brain activity better and more uniquely than the unimodal models. This is intriguing because human cognition is inherently multi-modal and multi-modal models might capture such a computational process in their latent representations. See Figure A.6 for the results for all subjects.

## 5 Discussion and conclusions

In this study, we quantitatively compared the relationship between different latent representations of LLMs and brain activity using diverse annotations of semantic content. To achieve this, we collected 8.3 hours of an fMRI dataset of brain activity recorded while participants watched extensively annotated videos. We demonstrated that the LLMs’ latent representations explain brain activity particularly well for high-level background story content compared with traditional language models. Moreover, our results show that different levels of semantic content are distributed differently in the brain. Finally, we demonstrated that multi-modal vision-semantic models explain brain activity better and more uniquely than unimodal models. While our findings align with prior work, we extend it by simultaneously providing a more detailed analysis of uniquely explained variances across multiple levels of semantic representations and brain regions, particularly when comparing multimodal and unimodal models. Overall, our findings provide valuable insights into interpreting LLMs from a neuro-AI perspective, offering a finer-grained understanding of the similarities between human cognition and AI compared to previous studies.

We reemphasize that these insights were not apparent from the modeling of individual features, as was performed in previous studies. For instance, the absence of unique variance for *Summary* and *TimePlace* is a significant insight, which indicates that simply including these types of information in encoding analysis might not be sufficient for capturing high-level semantic representations in the brain. Our results do not indicate that such information is absent in the brain but the absence could be caused by the limitations of the modeling approaches or fMRI measurement. Hence, in future work, we need to consider how to model these types of high-level information using our data as a new biological benchmark for alignment between LLMs and humans.

## 6 Limitations

Firstly, we constructed our encoding models using multiple (five) levels of semantic content. Although this approach is more comprehensive compared to previous research, which typically used only one level of semantic content, our study still does not encompass the full diversity of semantic content processed in reality. Moving forward, it will be crucial to use resources like LLMs to annotate stimuli and model a richer human semantic experience.

In this study, our focus was on the individual level, which is the most typical approach in the field of brain encoding models. We did not extensively compare how the representations of semantic content differ or shared among individuals. Since our dataset includes six individuals viewing the same stimuli, it will be crucial to compare these results in future research.

In this study, we used vision-language models to examine the importance of multi-modal fea-tures. However, human perception encompasses a broader range of modalities, including not just vision and language, but also hearing and other senses. To comprehensively understand these processes, it is important for future research to utilize models that can handle a greater variety of modalities.

In this study, while we used LLMs (e.g. Llama2) and multi-modal LLMs (e.g. Llava-v1.5), which were commonly used at the time of the research, more powerful models has been introduced continuously. It is important to examine whether the accuracy will improve further by using these more powerful models.

The overall accuracy for *Summary* and *Time-Place* were low, even when using LLMs. We believe this is not because LLMs do not correspond with the human brain in terms of these annotations, but rather due to potential issues with the modeling approach. In future studies, it will be necessary to develop methods that can more precisely capture such high-level information.

## Acknowledgements

Y.T. was supported by JST, PRESTO Grant Number JPMJPR23I6. SN was supported by MEXT/JSPSKAKENHI JP18H05522 and JP24H00619, as well as JST JPMJCR18A5 and JPMJCR24U2.

## Author Contributions

S.N. and Y.T conceived the study. N.K., H.Q.Y., R.K. and S.N. designed and performed the experiments. N.K., H.Q.Y., Y.N., T.M., and Y.T. analyzed the data. Y.T. wrote the original draft. S.N. and Y.T. wrote the manuscript with the consultation by the other authors.

## A Appendix

### A.1 fMRI dataset and preprocessing

#### A.1.1 Stimuli

In our study, we used multi-modal stimuli from nine DVDs, which encompassed 10 episodes of television drama series and a feature film, as detailed in Table A.2. The selection of these nine DVDs adhered to specific criteria: We chose 1) internationally acclaimed dramas and films, premised on the belief that their fame ensured the compelling nature of their content. This was intended to captivate the participants’ attention during the narratives in our study. 2) We included a diverse range of genres. Typically, a single story predominantly features certain characters whose dialogue and actions reflect the genre. In such limited scenarios, when deploying algorithms based on machine learning to analyze BOLD signals, ensuring the generalizability of results may prove to be challenging. To cover a broad spectrum of scenarios, we included films from multiple genres; The average total duration for the 10 episodes was 49.98 minutes (Table A.2). Each episode was segmented into two to nine segments for use in our imaging sessions. We chose the segmentation points to maintain segment lengths of approximately 10 minutes and to coincide with transitions between narrative scenes, thereby facilitating the participants’ comprehension of each episode. For each segment, except the initial segment, we included the concluding 20 seconds of the preceding segment. The resulting segments varied in length from 512 seconds to 1271 seconds, with an average of 746.8 seconds (Table A.2). Instead of converting each segment into a separate film file, we designated specific playback intervals to the respective DVDs using the intervals as visual stimuli in each imaging session. All films, with the sole exception of “Ghost in the Shell” (originally produced in Japanese), were presented in the Japanese-dubbed version, considering that all participants in our study were Japanese.

#### A.1.2 Procedures

fMRI BOLD signals were recorded as participants viewed the audiovisual content (the film segments) from 10 episodes across nine DVDs. The visual stimuli were projected centrally with a visual angle of 26.78 × 15.85 degrees at 25 or 30 Hz. MR-compatible headphones delivered the auditory stimuli. Prior to the viewing sessions, the audio levels were calibrated using non-experimental test clips to ensure clarity and a comfortable volume for participants. Participants were instructed to view the film segments casually, mirroring their everyday television-watching experience. For each participant, fMRI data were acquired over 10 distinct sessions. During each session, participants watched one or two episode segments over three to five segments (each segment lasting approximately 10 minutes) (Table A.2). Because of its extended length (about 2 hours), “Dream Girls” was viewed over two sessions. Each segment’s film content was displayed by cueing the respective DVD to play at specific times using the VLC media player’s (VideoLAN, France) command line interface. This interface was set up to commence playback coinciding with the scan’s start. Manual termination of the scan followed the film segment’s conclusion, thereby accommodating the varying durations of playback across segments and sessions. In total, around 9 hours (31905 seconds) of film content were presented to the participants across 10 sessions for fMRI data collection.

#### A.1.3 Preprocessing

For individual preprocessing of EPI data for each participant, the Statistical Parameter Mapping toolbox (SPM8, http://www.fil.ion.ucl.ac.uk/spm/software/spm8/) was used. EPI images were motion-corrected by aligning them to the initial image recorded in the first session for each participant. Voxel responses were standardized by deducting the average response over all time points. Subsequently, prolonged trends in the standardized responses were mitigated by detracting the outcome of median filter convolution with a 120-second time frame. Data standardization and detrending were conducted for each movie segment for each voxel. Data captured within the initial 20 seconds of the scan were deemed susceptible to artifacts from startup transients and thus excluded from the analysis. The analysis considered data from 20 seconds post-scan initiation to the conclusion of the film content.). To account for the hemodynamic response function (HRF), stimulus features were time-shifted by 8s and then averaged with the stimulus feature corresponding to each volume and the feature corresponding to the subsequent two-second volume. See Figure A.7

#### A.1.4 Interpretation of PCA

In the annotation process using GPT-4, we allowed the annotations to be associated with multiple semantic attributes. Then, we evaluated each PC according to the average scores for the five attributes. The five attributes depict a tense or peaceful environment (“Tense confrontations or crime” and “Everyday interaction or peaceful living”), represent individual or collective decision-making (“Personal growth, change, or determination”, and “Decision-making or role of leaders”), and illustrate interaction or cooperation with others (“Mutual assistance or cooperation’). We also define these five attributes using GPT-4 by asking GPT-4 to identify common attributes across annotations.

To further interpret information content in the PCs, we projected *Story* annotations on each PC. Again, we observed consistent trends in PC1 and PC2. Figure 4b shows that PC1 contrasts annotations related to individual or collective decision-making (“Decision-making or role of leaders” and “Personal growth, change, or determination”) with annotations depicting environment or background scenario (“Tense confrontations or crime” and “Everyday interaction or peaceful living”). Regarding PC2, it contrasts annotations that indicate cooperation with others (“Decision-making or role of leaders” and “Mutual assistance or cooperation”) with more personal scenarios (“Decision-making or role of leaders”, “Everyday interaction or peaceful living,” and “Tense confrontations or crime”). Regarding PC3, although there was a tendency among participants to contrast cooperation (“Mutual assistance or cooperation”) with other attributes, the variation across participants was large.

#### A.1.5 Feature extraction

We use bert-base-uncased, gpt2-large, facebook/opt-6.7bm, meta-llama/Llama-2-7b-hf, MIT/ast-finetuned-audioset-10-10-0.4593, facebook/mms-1b-models, facebook/deit-base-distilled-patch16-224, microsoft/resnet-50, microsoft/git-base, BridgeTower/bridgetower-base, and llava-hf/llava-1.5-7b-hf models available on Hugging Face for BERT, GPT2, OPT, Llama2, AST, MMS, DeiT, ResNet, GIT, BridgeTower, and Llava-v1.5. We use GoogleNews-vectors-negative300 model available on https://code.google.com/archive/p/word2vec/.

### A.2 Annotation procedure

Each video was annotated for five types of semantic content, including utterance sounds, context, and background stories. These annotations were selected because they are critical for narrative understanding. While other annotations, such as discourse structure, dialogue acts, and various syntactic and semantic analyses, are useful, our study specifically aims at the natural flow and comprehension of narratives.

The annotations were performed by one or several annotators, employed by external agencies, for each type of semantic content.

Note that *Speech* annotations refer to the exact content spoken by actors in a video (e.g., “Hey John, how are you feeling?” “Great”). These can indeed be expressed as language descriptions. On the other hand, *Story* pertains not to the direct dialogue but to the context or background information (e.g., “Mary and John, who have been childhood friends, are reuniting after two years”). This allows it to be described as a separate linguistic content from the spoken dialogue.

The overview of each annotation and attributes of the annotators are as follows. The details of the annotation procedures will be more thoroughly explained in the documentation accompanying the future release of the dataset.

1. *Speech*: Describe the speech and narration content, specifying the start and end times of each segment. A single male annotator in his 40s annotated all the content.
2. *Object*: Describe the content displayed on the screen every second. The descriptions should be in natural language and range from one to several sentences. Thirteen annotators annotated the content, with 3 in their 30s, 7 in their 40s, and 1 in 50s. Eleven of the annotators were female.
3. *Story*: Approximately every five seconds, describe the story based on the content of the narrative. Three annotators annotated the content, with 1 in 30s and 2 in their 40s. Two of the annotators were female.
4. *Summary*: Approximately every 1–3 minutes, or at points where the overall flow of the story changes, describe the content of each section and provide summaries of the actions performed by each character. Eight annotators annotated the content, with 7 in their 30s and 1 in 40s. Six of the annotators were female.
5. *TimePlace*: At each scene transition, describe the time of day and location. Three annotators annotated the content, with 1 in 30s and 2 in their 40s. Two of the annotators were female.

Note that, for *Speech* annotations, we transcribed the words actually spoken in the video. Specifically, we collected speech transcription from two annotators. One of the annotators was an English speaker and transcribed English speech while watching the movie with English audio. The other was a Japanese speaker and transcribed Japanese speech while watching a Japanese-dubbed version of the movie. For each language, we split the speech into units as short as possible to maintain the meaning of sentences, and annotated the transcription and the onset/offset of each speech part. The transcription included speech, filler, and non-verbal utterances (e.g. laughing voice and a cough) attributed to each character. Note that English and Japanese speeche coincided in both the timing of delivery and the intended meaning, but they were not entirely congruent.

Below are three example annotations from three different scenes in our dataset. For *Summary*, the description of the content section is extracted and displayed:

- Scene from *Suits*:
  – (*Speech*) *A novel? A grade schooler?*
  – (*Object*) *The face of a man in a suit is shown large in the center of the screen. The man looks forward, mouth closed, and appears to have a serious expression on his face*.
  – (*Story*) *Harvey expressed surprise that Mike, an elementary school student, was reading an adult novel. Mike replied that he loved books*.
  – (*Summary*) *Harvey Spector is interviewing Mike Ross. He is interested in Mike Ross when he hears how he was able to identify him as a police officer. He gives Mike Ross a question to test his knowledge. Harvey Spector notices that Mike Ross has a good memory and is smart, and decides to hire Mike Ross as an associate*.
  – (*TimePlace*) *Noon. Hotel. Room*.
- Scene from *The Crown*:
  – (*Speech*) *It’s okay. It’s all right*.
  – (*Object*) *A man is shown in the center looking to the left. Behind the man is a stained glass window, and several figures are vaguely visible*.
  – (*Story*) *Philip was waiting for the woman (Elizabeth) in the church with a nervous look on his face. He told himself over and over again that he would be fine*.
  – (*Summary*) *In front of the church altar, Philip looks nervous as he waits for the ceremony to begin. Former Prime Minister Winston Churchill and his wife arrive at the church, where the attendees stand to greet them*.
  – (*TimePlace*) *Morning. Chapel*.
- Scene from *Ghost in the Shell: STAND ALONE COMPLEX*:
  – (*Speech*) *Hey, what’s going on?*
  – (*Object*) *Two men are wearing uniforms and have their hands clasped behind their backs, as if guarding a figure sitting in a chair. Behind them, a sharp-eyed man appears to be on the phone*..
  – (*Story*) *At air traffic control, one man with glasses is impatient and tries to figure out what’s going on by whispering on the phone*.
  – (*Summary*) *A tank suddenly starts moving inside the experimental facility of Kenryo Heavy Industries. When a worker calls out to it, it stops, turns around, and attacks the worker and other tanks with a machine gun and cannon. Unrest spreads among the workers and air traffic controllers. Ohba watches the scene from a short distance outside the fence*.
  – (*TimePlace*) *Morning. Inside the Dome. Management Office*.

### A.3 Quality control of the dataset

To verify the quality of the dataset, we split the dataset and annotations as follows to ensure that the results were consistent when the encoding model was built separately for each split.

First, we split the entire training dataset into two parts: runs of the first and second halves of the data used in the main analysis. We used the same videos for testing dataset. Regarding ‘Breaking Bad,’ because only one run of data was used in the main analysis, we excluded it from this control analysis. The comparison shows that the results for the two encoding models using different data splits produced quite similar results across cortical voxels (see Figure A.9).

We also checked the quality at the annotation level. For our data, there were two annotators for *Story* and five for *Object*. Therefore, for *Story*, we split the two annotators and these split are used to create their respective encoding models. For *Object*, we split the annotators into groups of three and two and used to create their respective encoding models to compare the results. The comparison shows that the results for the two encoding models using different annotator splits produced quite similar results across cortical voxels (see Figure A.10).

Finally, for the *Speech* annotations, we performed analysis to verify the consistency in the speech transcription between Japanese and English. Specifically, we compared the results for the encoding models when we used Japanese *Speech* annotation with the results when we used English *Speech* annotation with the latent representations of GPT2. In this analysis, we constructed the encoding model for the two languages, respectively, and compared the prediction accuracies across voxels for each participant. We then examined whether the prediction accuracies exhibited a similar pattern between two annotations. We observed strong correlation across all participants (DM01: Pearson’s r = 0.90, DM03: r = 0.85, DM06: r = 0.91, DM07: r = 0.90, DM09: r = 0.92, DM11: r = 0.82), which indicates that the latent features for the two languages had similar information to explain brain activity.

### A.4 Additional results of encoding models

Figures A.1, A.3, A.5, and A.6 show additional results for all subjects for Figures 2, 3, 4, and 5, respectively. They show that our results were robust across subjects.

**Table A.2:**
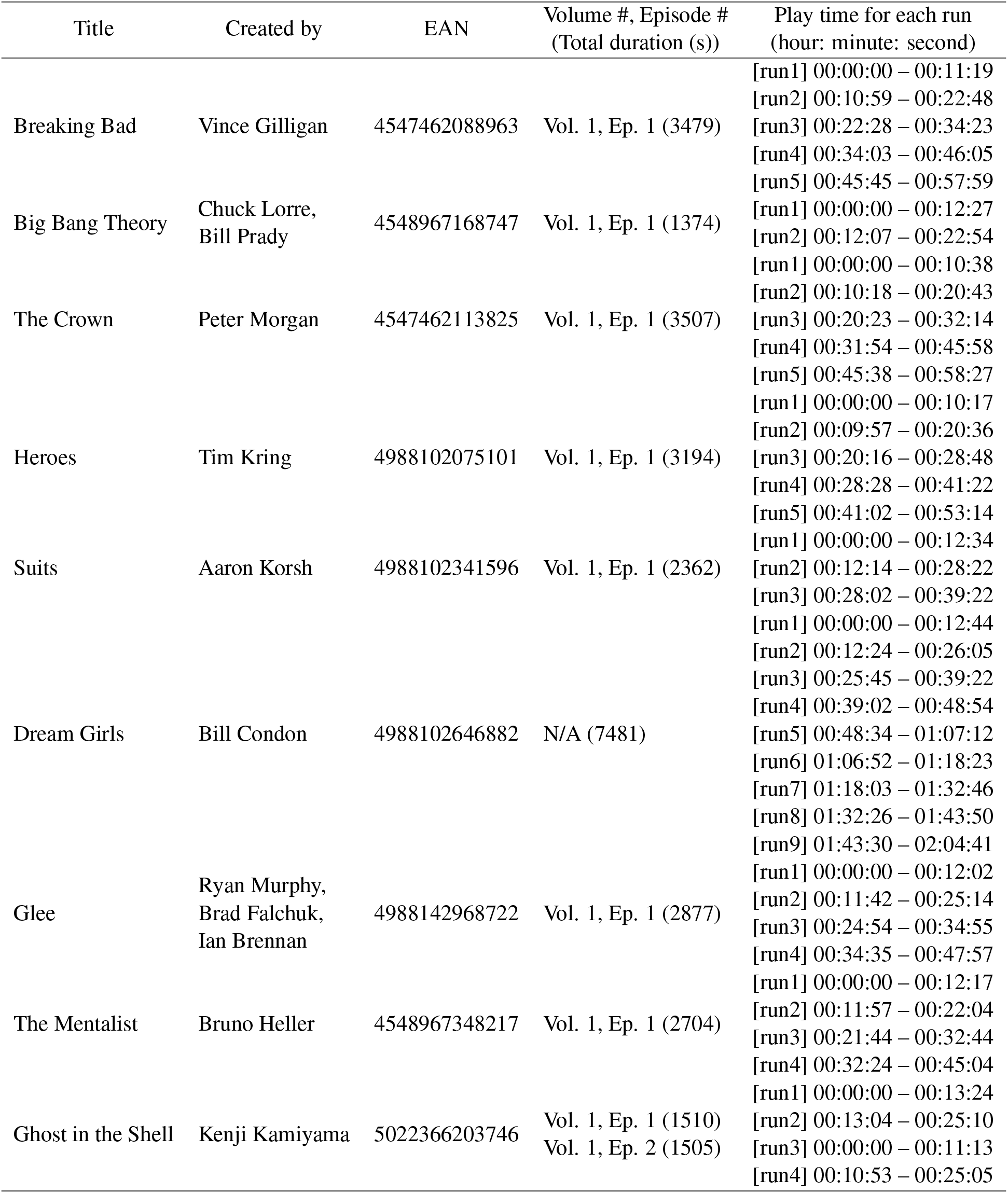
Videos used for the experiment.

**Figure A.1:**
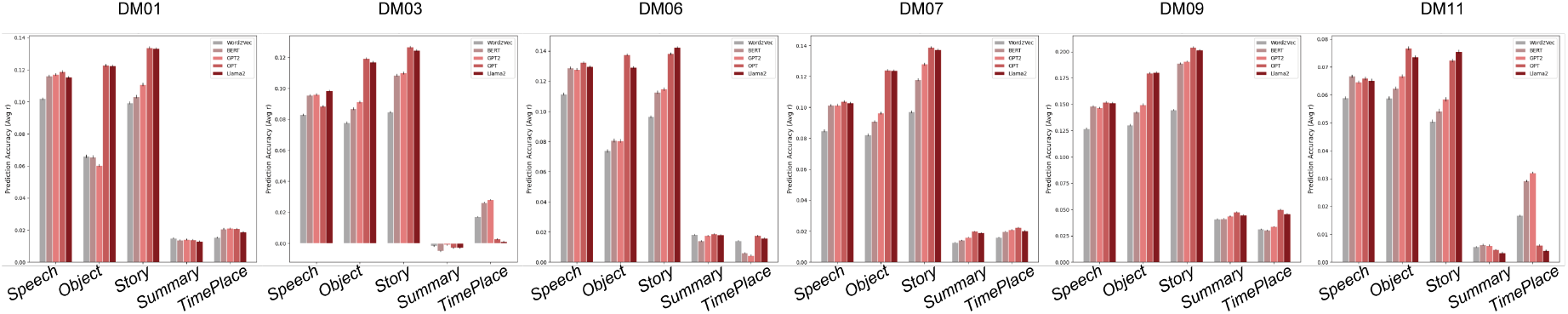
All subject results for comparing prediction performances among different language models for different semantic contents.

**Figure A.2:**
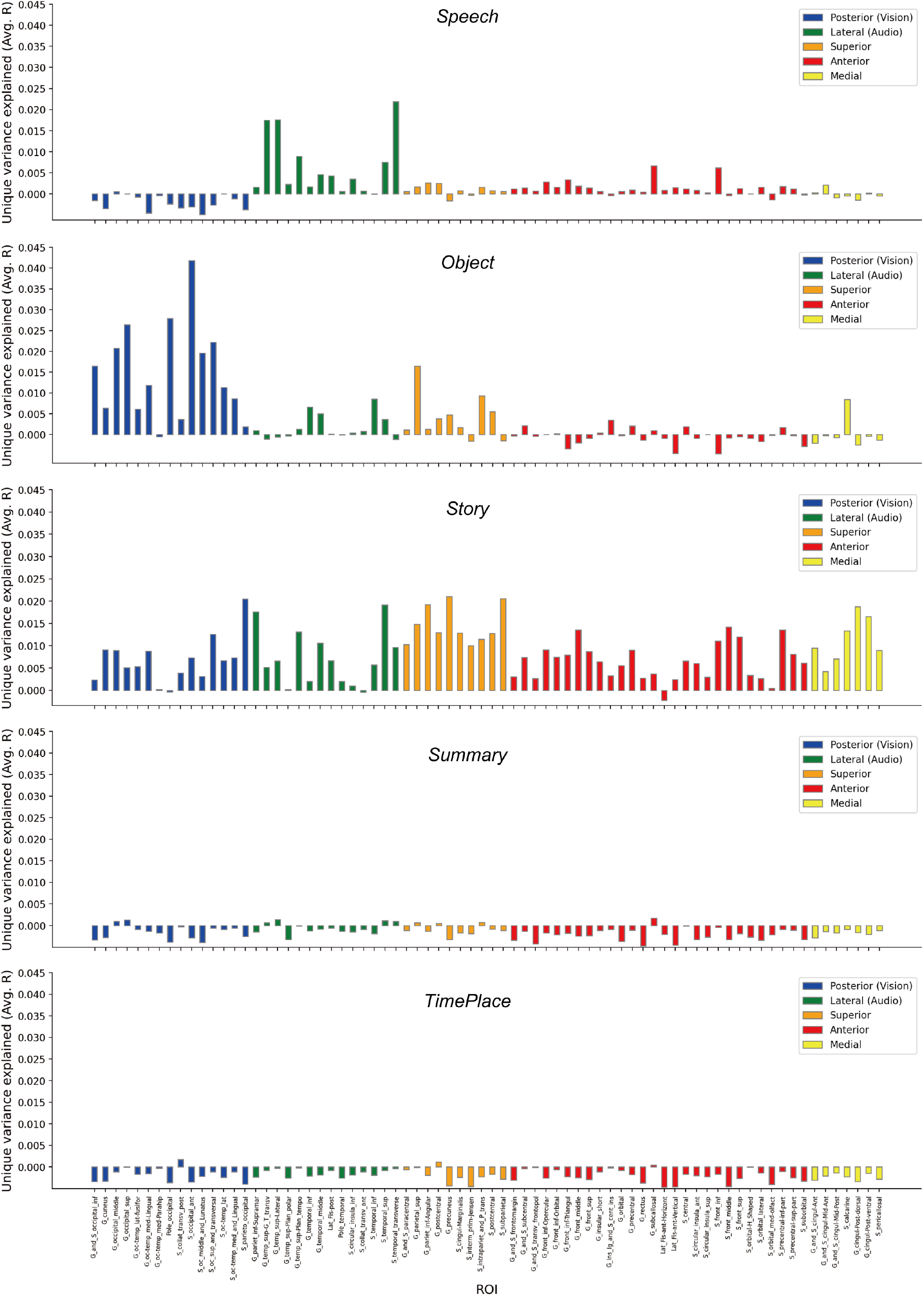
Single subject (DM09) result for comparing prediction performances among different ROIs for different semantic contents. Each bar represents averaged unique variance explained across voxels within each ROI

**Figure A.3:**
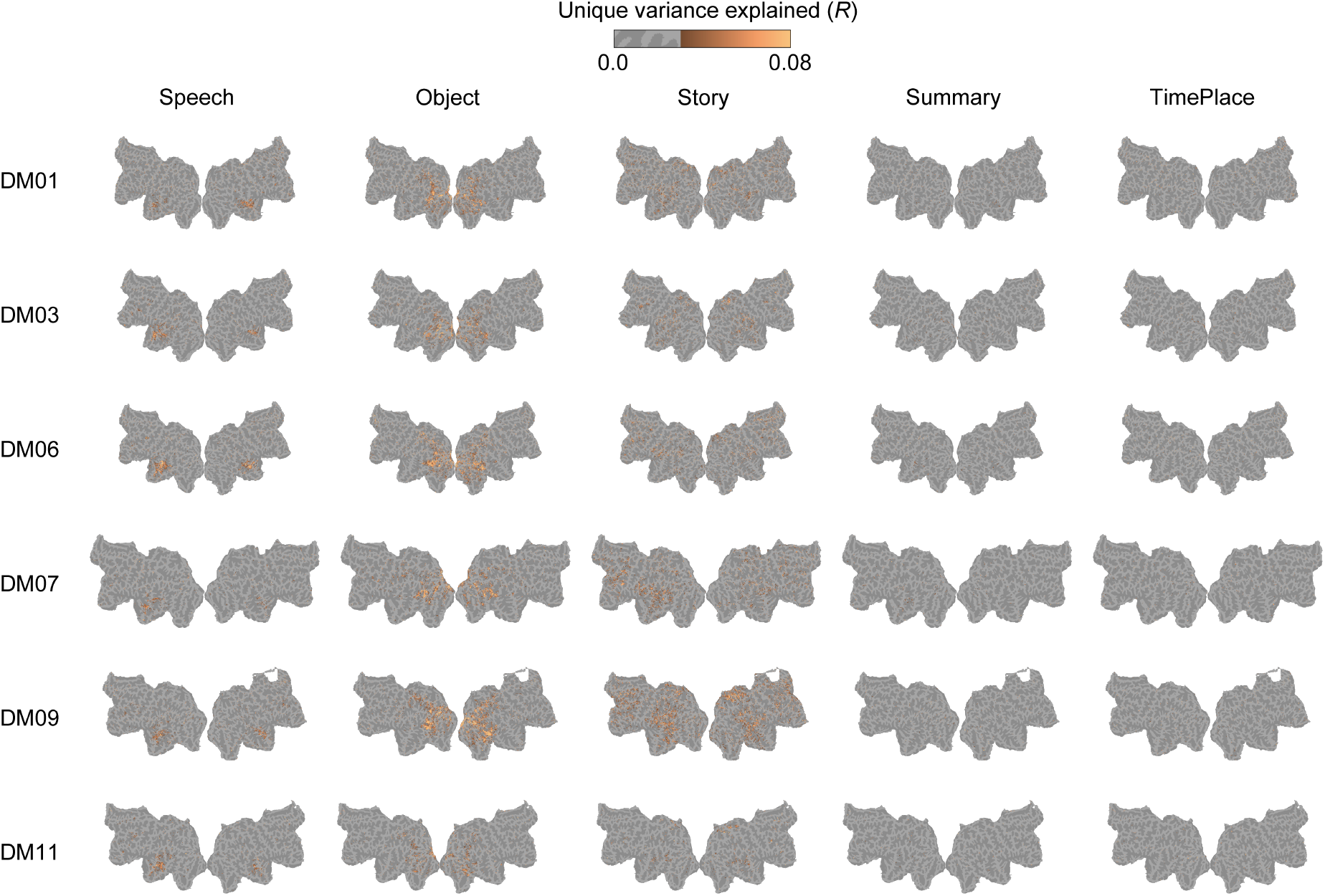
All subject results for comparing unique variance explained among different semantic contents using variance partitioning analysis.

**Figure A.4:**
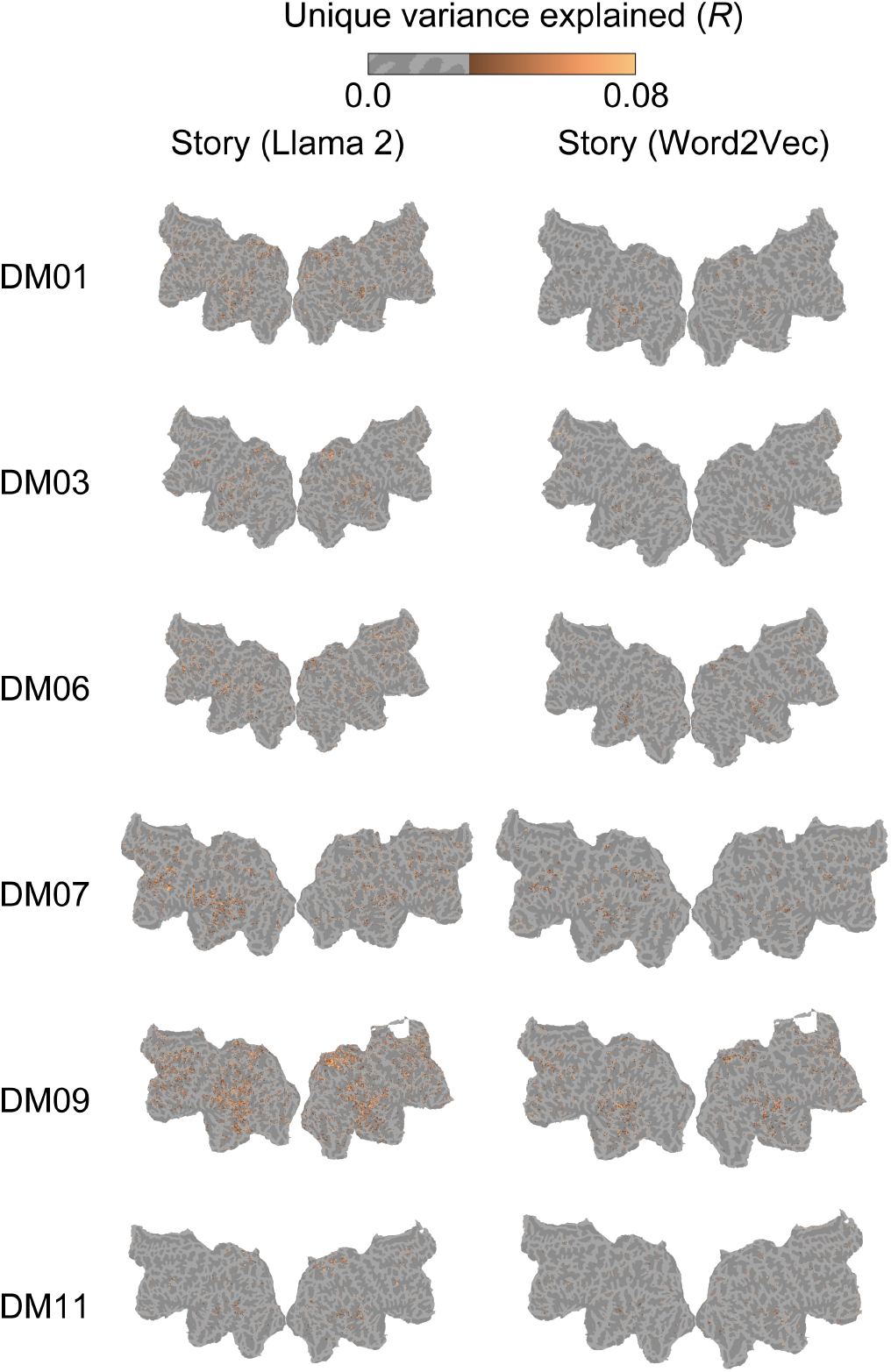
All subject results for unique variance explained of for *Story* feature for Llama 2 (Left) and Word2Vec (Right), respectively.

**Figure A.5:**
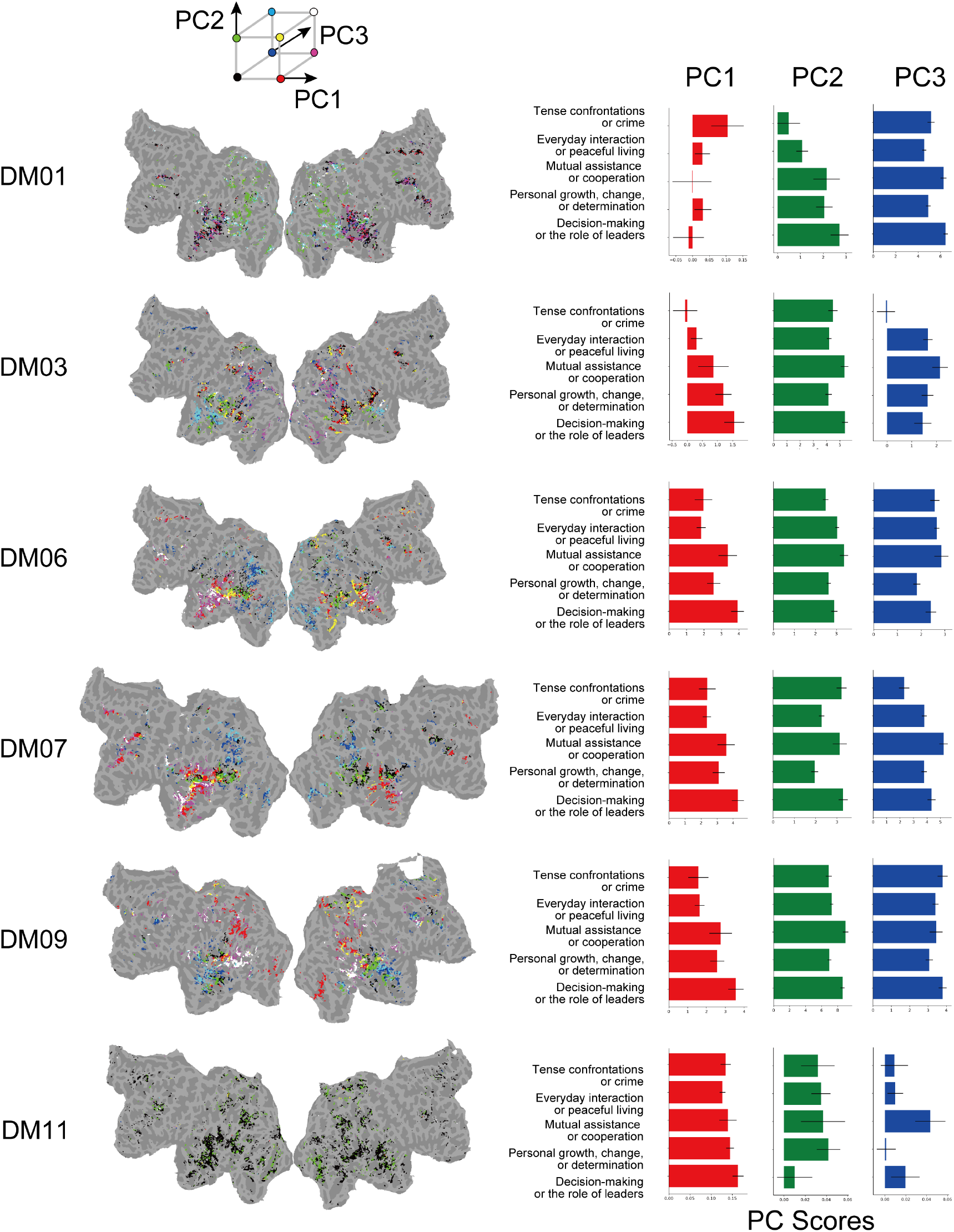
All subject results for principal components analysis. We project each caption onto the PC space and then calculated the PC scores for each attribute assigned to the caption. The error bars represent the standard error of the mean PC scores across annotations belonging to each attribute.

**Figure A.6:**
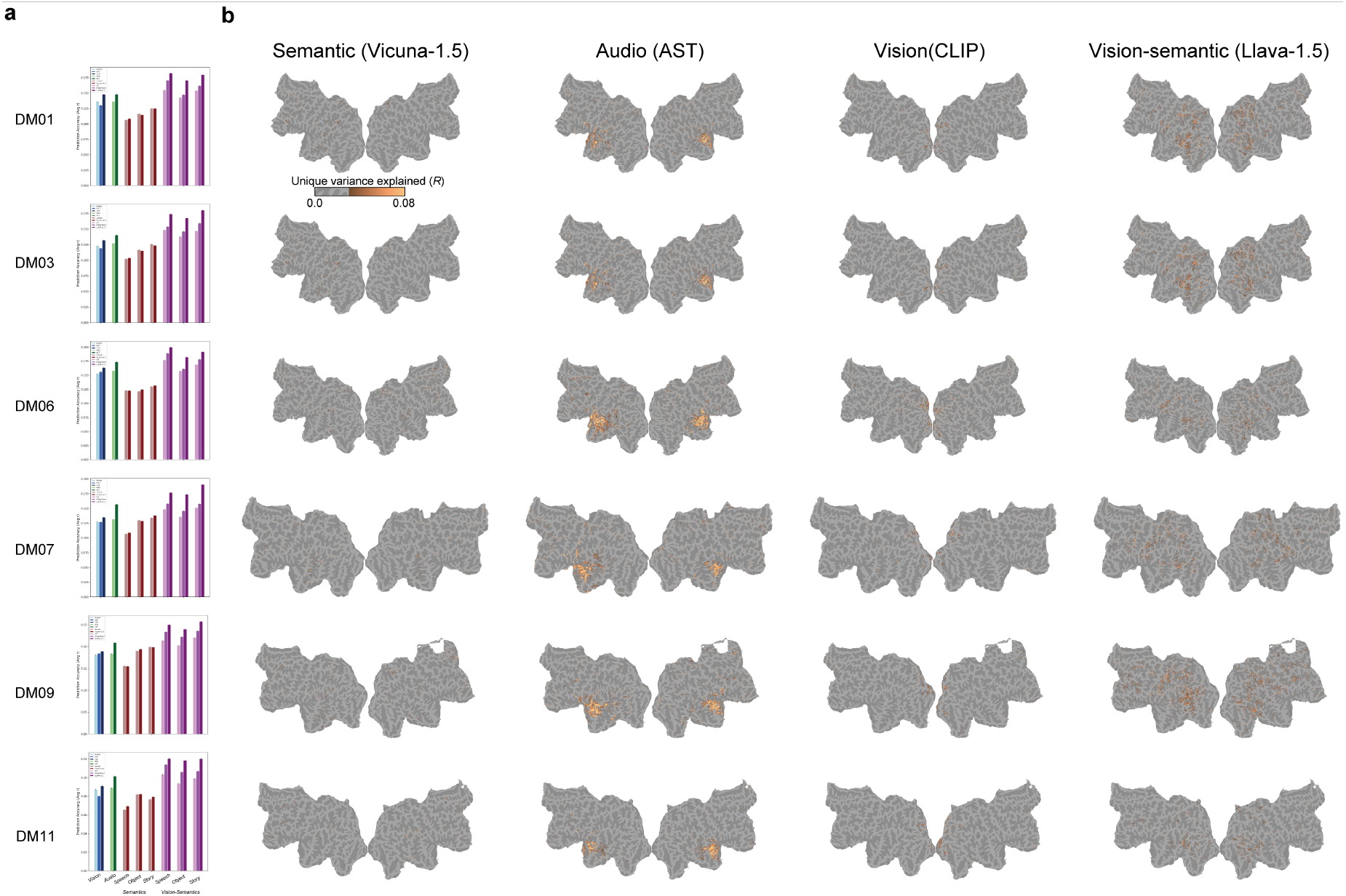
All subject results for comparing the prediction performance among semantics, audio, visual, and vision-semantic features.

**Figure A.7:**
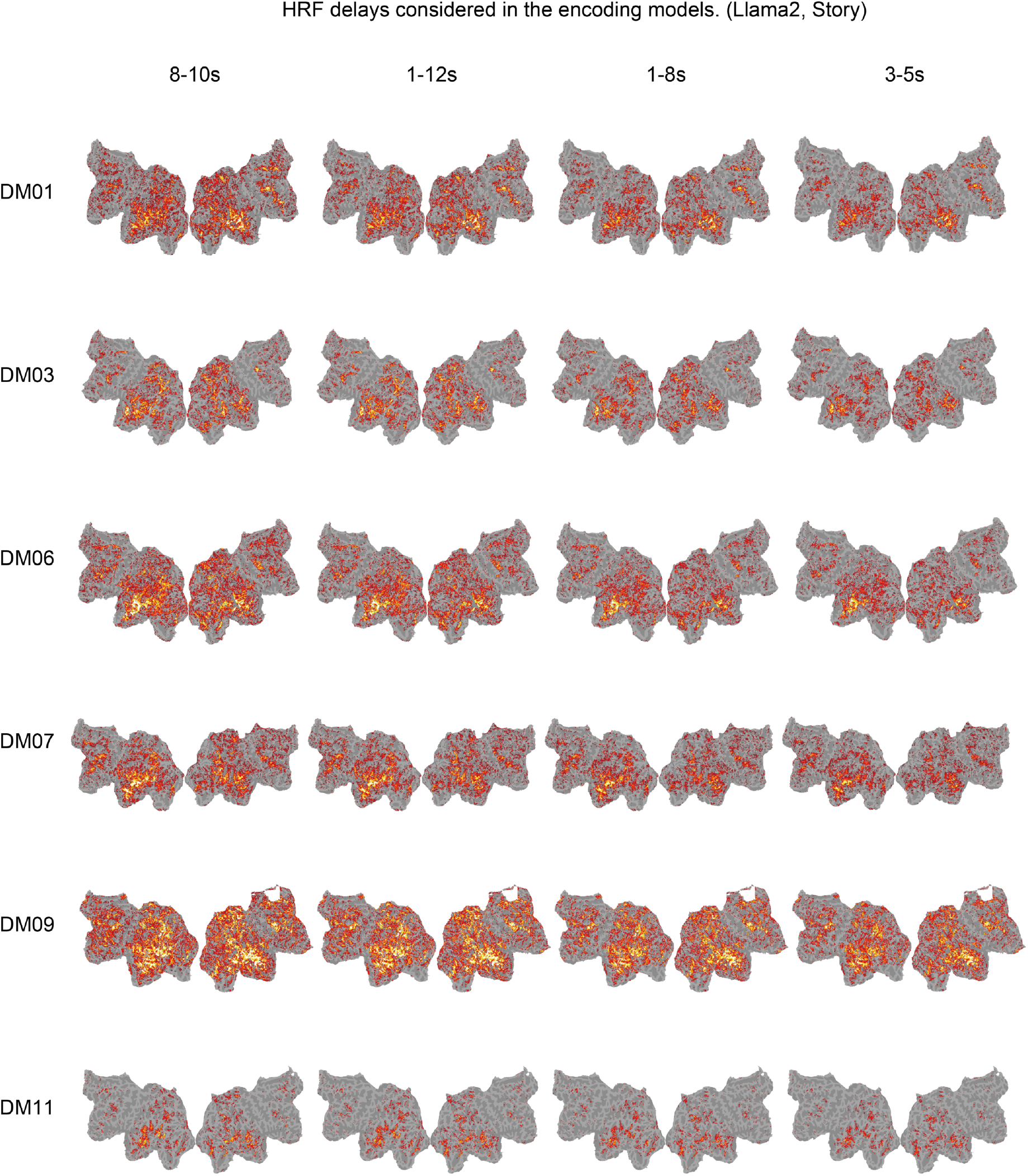
Results for the encoding models under the assumption of different BOLD signal time delays, using the Llama 2 with *Story* Feature. Changing the settings used in the main analysis (8-10s) does not affect the overall patterns of results.

**Figure A.8:**
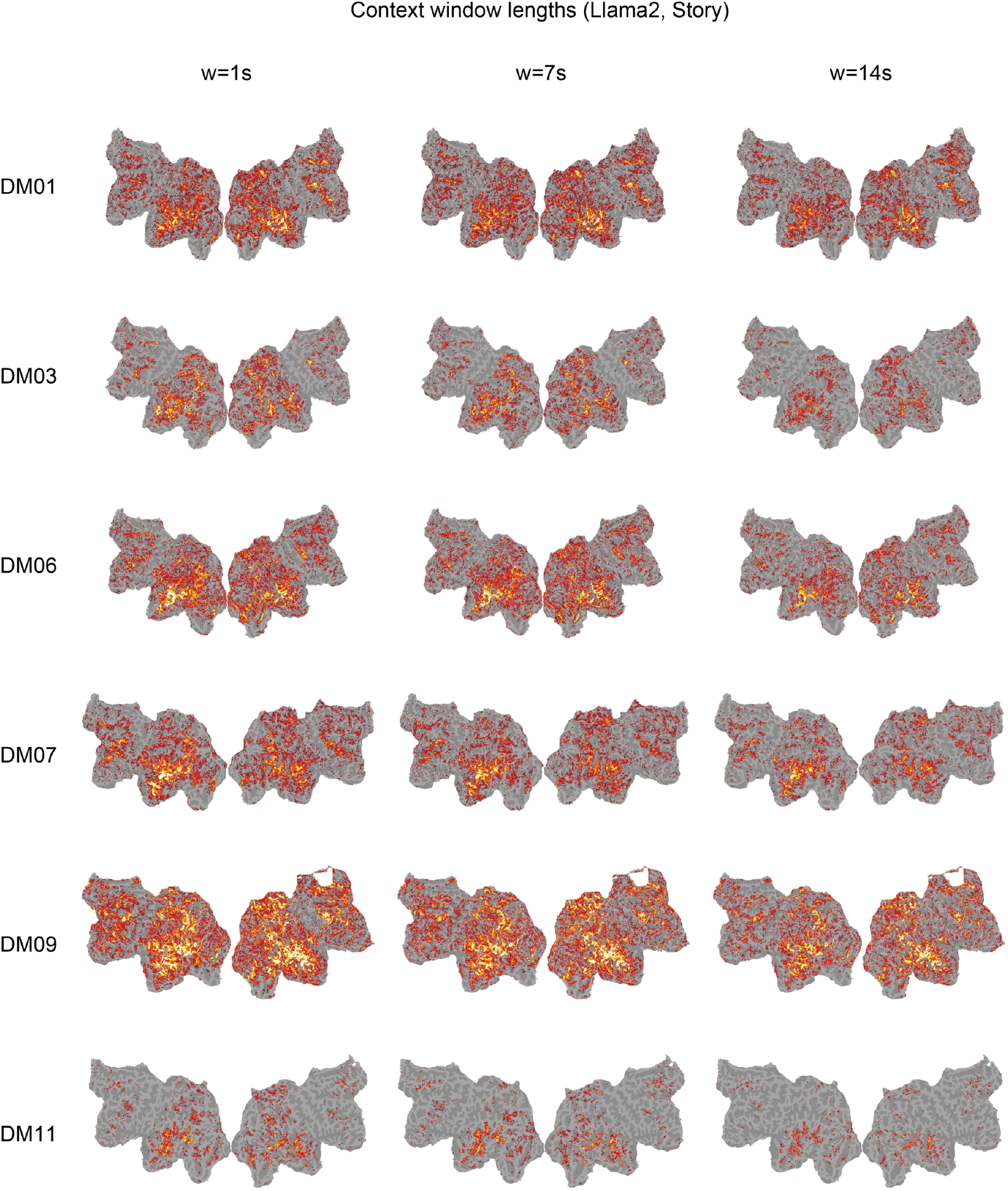
Result for the encoding models using the *Story* feature of Llama 2 with different context widths (w). Changing the setting used in the main analysis (w=1s) does not affect the overall patterns of results.

**Figure A.9:**
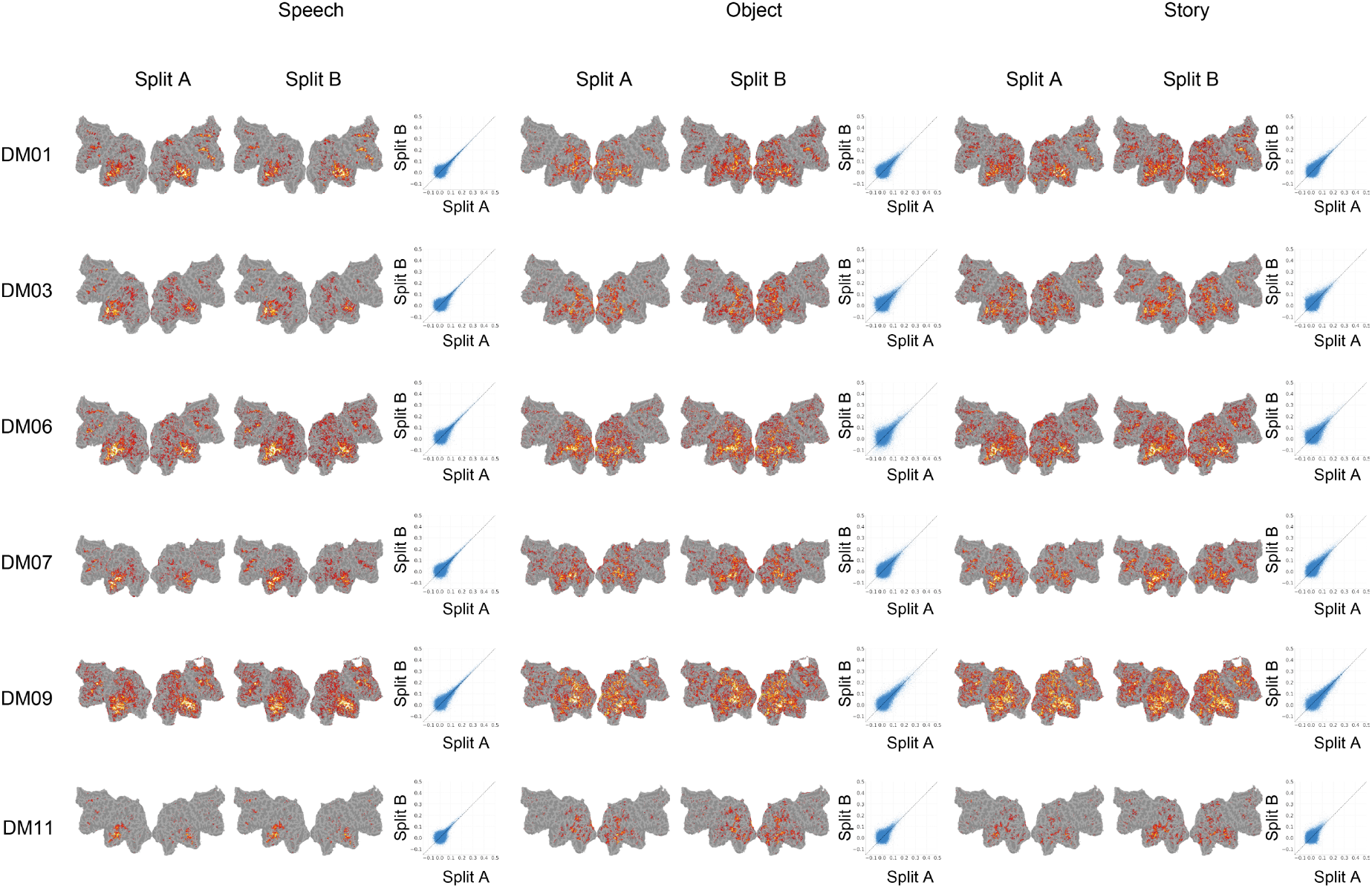
Results for encoding models when training data are split in half (Split A and Split B) and the models are built independently. We observe no significant difference in the distribution of the flat map in both splits. Also, when comparing prediction performance across voxels between Split A and Split B, the results are generally reproduced in both splits (see scatter plots).

**Figure A.10:**
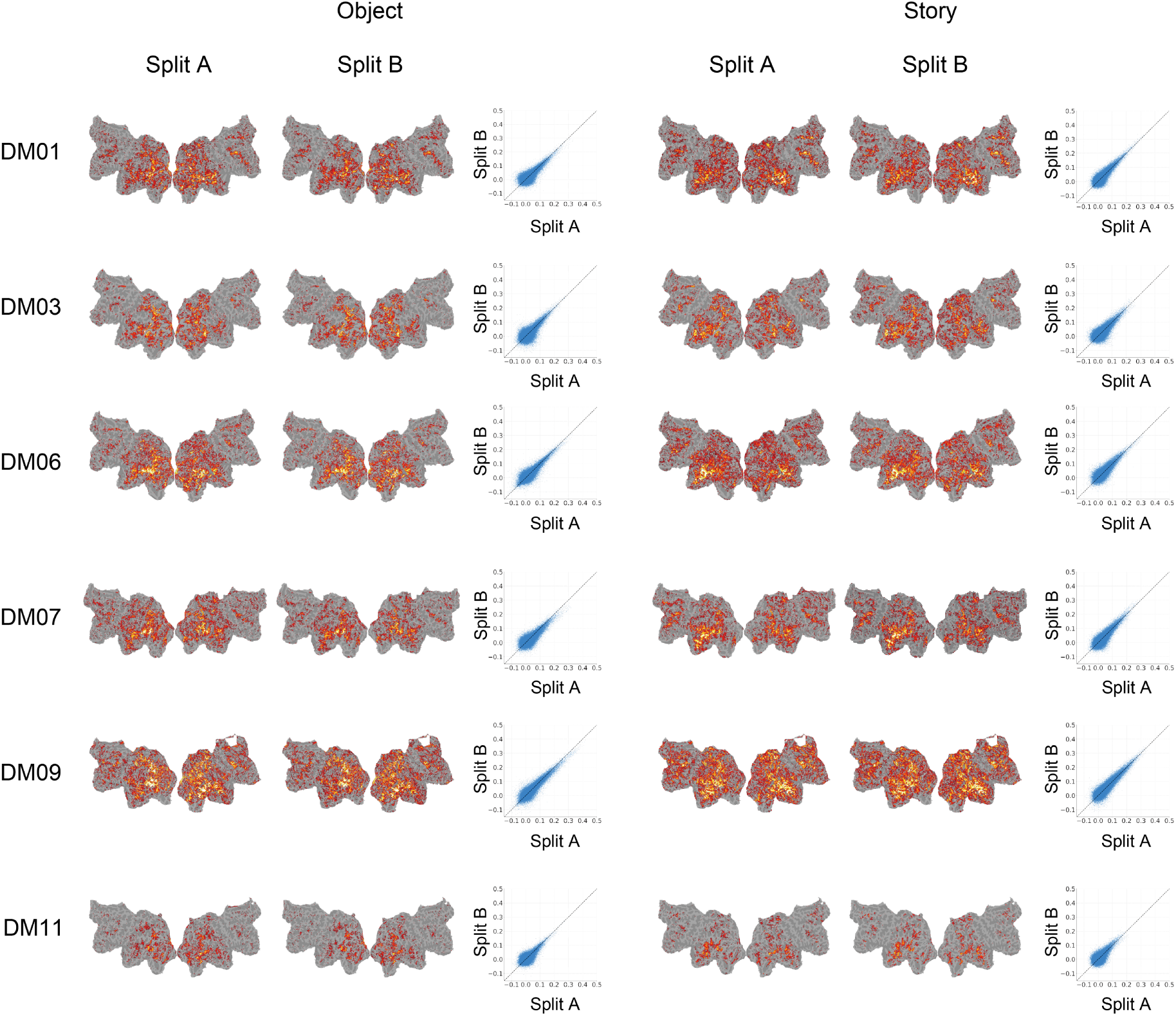
Results for encoding models when annotations are divided into two splits (Split A and Split B) and the models are built independently. We observe no significant difference in the distribution of the flat map in both splits. Also, when comparing prediction performance across voxels between Split A and Split B, the results are generally reproduced in both splits (see scatter plots).

## Footnotes

1 https://github.com/yu-takagi/drama2brain

2 We have confirmed that similar results are obtained by using LLMs trained in Japanese with original Japanese captions.

4 https://github.com/gallantlab/himalaya

